# Video tracking of single cells to identify clustering behaviour

**DOI:** 10.1101/2024.11.08.622624

**Authors:** Mónica Suárez Korsnes, Håkon André Ramberg, Kristin Austlid Taskén, Reinert Korsnes

## Abstract

Cancer cell clustering is a critical factor in metastasis, with cells often believed to migrate in groups as they establish themselves in new environments. This study presents preliminary findings from an *in vitro* experiment, suggesting that co-culturing cells provides an effective method for observing this phenomenon, even though the cells are grown as monolayers. We introduce a novel single-cell tracking approach based on graph theory to identify clusters in PC3 cells cultivated in both monoculture and co-culture with PC12 cells, using 66-hour time-lapse recordings. The initial step consists of defining ‘linked’ pairs of PC3 cells, laying the foundation for the application of graph theory. We propose two alternative definitions for cell pairings. The first method, *Method 1*, defines cells as ‘linked’ at a given time *t* if they are close together within a defined time period before and after *t*. A second potential alternative method, *Method 2*, pairs cells if there is an overlap between the convex hulls of their respective tracks during this time period. Pairing cells enables the application of graph theory for subsequent analysis. This framework represents a cell as a vertex (node) and a relation between two cells as an edge. An interconnected set of high-degree nodes (nodes with many connections or edges) forms a subgraph, or backbone, that defines a patch (cluster) of cells. All nodes connected to this backbone are part of the subgraph. The backbone of high-degree nodes functions as a partition (or cut) of the initial graph. Two consecutive clusters in the video are considered to share the same identity if the following cluster contains at least *p* = 75 % of the cells from the preceding cluster, and the mean positions of their cells are within △*r* = 75 µm. PC3 cells grown in co-culture appear to form persistent clusters exceeding 10 cells after 40 to 50 h incubation following seeding. In contrast, PC3 cells cultured alone (mono-culture) did not exhibit this behaviour. This approach is experimental and requires further validation with a broader dataset.

## 1 Introduction

This work aims to improve access to data on clustering within cancer cell populations. A cluster is defined as a small group of cells that maintain spatial proximity and exhibit coordinated behaviour over time [Gopinathan and Gov, 2019]. This phenomenon is thought to play a critical role in metastasis, and new observations could enhance our understanding and lead to novel approaches to cancer treatment. Understanding the characteristics and dynamics of these clusters is essential for uncovering the mechanisms behind cancer progression, and single-cell clustering assays offer a powerful approach to studying how such cell groups form, interact, and evolve over time.

Single-cell clustering assays can reveal how cells communicate and influence each other, particularly when they cluster and respond collectively. Analysing clustering can help identify early patterns of abnormal cell behaviour, potentially leading to earlier disease diagnosis. Several authors have contributed to advancing this field [Liu et al., 2023, Maity et al., 2024, Shannon et al., 2024].

The present preliminary *in vitro* experiments suggest that co-culturing cells may promote increased clustering. This finding stems from a clustering analysis approach inspired by graph theory [Ioannides et al., 2022]. Our method utilizes cell movement data to identify clusters, as demonstrated by analysing 66-hour time-lapse recordings of PC3 prostate cancer cells in both monoculture and co-culture with PC12 nerve cells. These data suggest that PC3 cells interact in ways that influence their movements, particularly when co-cultured with PC12 cells. Lasting clusters begin to form after approximately 35 hours of recording when PC3 cells are co-cultured with PC12 cells.

The current focus on using monolayer (2D) cells can, in principle, be generalized to apply to analyse recordings of cells grown in 3D, which offer an environment more similar to in vivo situations. However, it is advantageous to first test the 2D approach, as it can more easily provide tracking data over several generations. Recent advances in deep learning-based cell tracking may change this situation [Freckmann et al., 2022, Merino-Casallo et al., 2022, Mosier et al., 2021, Wen and Kimura, 2022, Wiggins et al., 2023].

The rationale for this showcase of methods is the general idea that unicellular organisms can exhibit collective behaviour, such as flocking [Ling et al., 2019]. Bacterial biofilms, where bacteria work together to form complex structures, are well-established examples of how unicellular organisms can enhance their survival and proliferation. Cells are in general competent to produce a quorum signal [Niu and Wang, 2012]. Cooperation is useful for individuals to reach a collective benefit, share information or neutralize threats [Wrenn et al., 2021].

Cancer cells are often associated with selfish behaviour. However, they can cooperate and maintain physical contacts by forming clusters. This may facilitate the metastatic cascade and promote disease progression [Archetti and Pienta, 2019]. Such clusters of cells appear to include distinct cellular states/phenotypes in which “leaders” and “followers” can affect the migratory pattern of clusters, as observed in melanoma and breast cancer Haeger et al. [2020], Khalil et al. [2017]. Such “leaders” and “followers” need to sense the attractant through the extracellular matrix integrin signalling and adhere to each other to coordinate their movements with robust directionality [Colak-Champollion et al., 2019].

A leader-follower organization among cells is important for successful invasion and metastasis [Wrenn et al., 2021]. Disrupting this organization can therefore greatly suppress the collective migration and metastatic potential [Cheung et al., 2013, Gao et al., 2017, Khalil et al., 2020, Yang et al., 2019, Zhang et al., 2019]. A recent study by Gómez-de Mariscal et al. [2024] demonstrated collective cell behaviour over a 14-hour period, with images captured at 10-minute intervals. They showed, for example, that cells closer to the leading edge exhibited more directional movement compared to those farther away. While the distance of cells to the leading edge remained constant from the beginning to the end of the tracking period, the distribution of the data was broad.

Co-culture experimental systems to identify cooperative phenotypes between “leaders” and “followers” have shown that leader cells can maintain their invasive phenotype and that the traditional EMT signature alone cannot be utilized to identify them [Konen et al., 2017]. Differences in metabolism, epigenetic modifications and gene mutations might also be important for identification of cooperative behaviour [Commander et al., 2020, Summerbell et al., 2020, Zoeller et al., 2019].

The outline of this paper is as follows. Sections 2.2–2.4 detail the production of test video of PC3 cells in mono-culture as well as in co-culture with PC12 cells. Sections 2.5.1-2.5.2 describes refinement of cell positional data obtained from singe-cell tracking, as well as demonstrating how to distinguish between PC3 and PC12 cells in video based on a nearly simultaneous fluorescence image at start. Section 2.5.3 provides two alternative methods to identify patches among PC3 cells based on track data from them. Section 3.1 brings a comparison between these two methods, suggesting that the choice of method may not be critical for identification of patches. Section 3.2 shows an example where PC3 cells in co-culture seem to form more stable clusters as compared to when they live in mono-culture. Section 4 discusses the limitations of the present method showcase, which is based on limited test data.

## 2 Materials and methods

### 2.1 General work flow

Our approach for cluster identification involves the following steps:

1. Cultivation of PC3 and PC12 cells (Section 2.2).
2. Capture a still image of the cells (with PC3 cells labelled with GFP) and record a video of them in monoculture and co-culture with PC12 cells (Figure 2.4.
3. Identify PC3 and PC12 cells in the co-culture by comparing the still image to the video image nearest in time (see Section 2.5.2 and Figure 1).
4. Track individual cells, ensuring that PC3 cells maintain their GFP label throughout their lineage. Refine and manually correct the tracks as needed (Section 2.5.1).
5. Identify clusters based on the refined tracks (Section 2.5.3).

**Figure 1.**
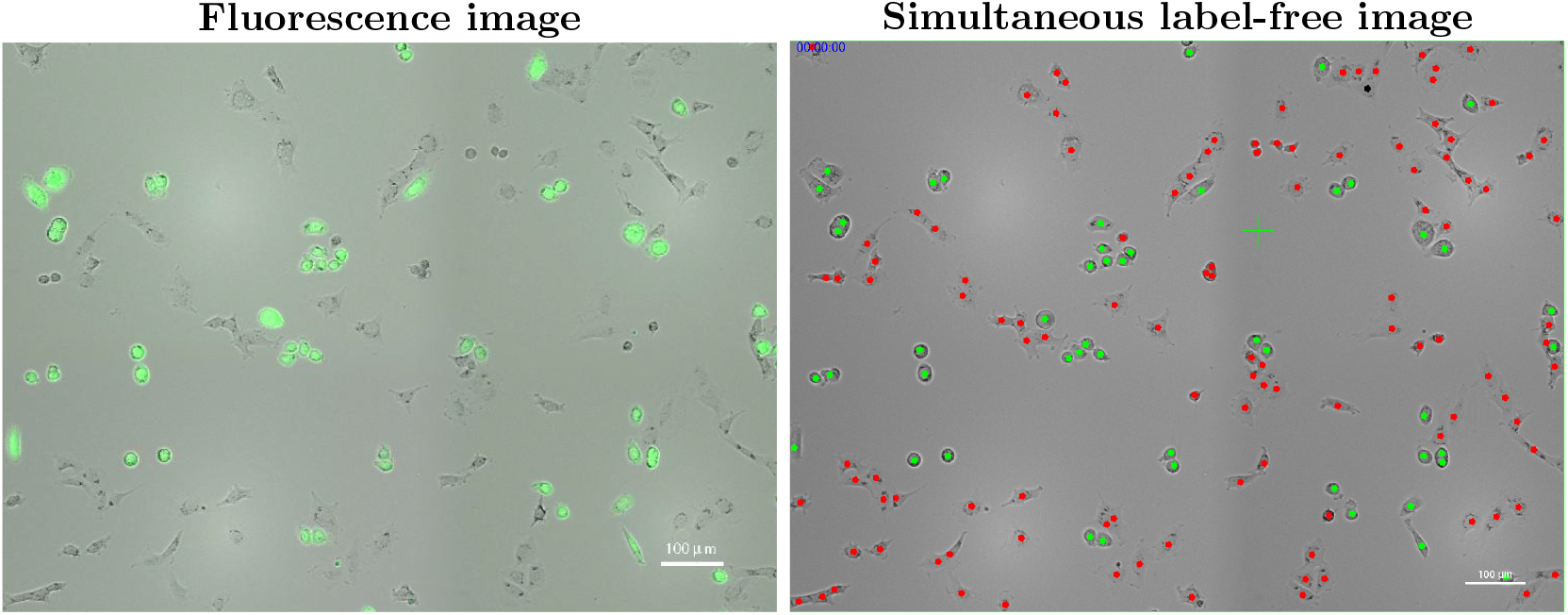
Manual classification of cells in a label-free video image (right, at the start of recording) using nearly simultaneous GFP fluorescence imagery (left), where GFP labels PC3 cells with green fluorescence. Green dots in the label-free video image correspond to GFP-labeled PC3 cells, while red dots indicate classified PC12 cells. Classifying a single cell in the video image based on its GFP fluorescence (green) allows for the classification of all members of its lineage.

**Figure 2.**
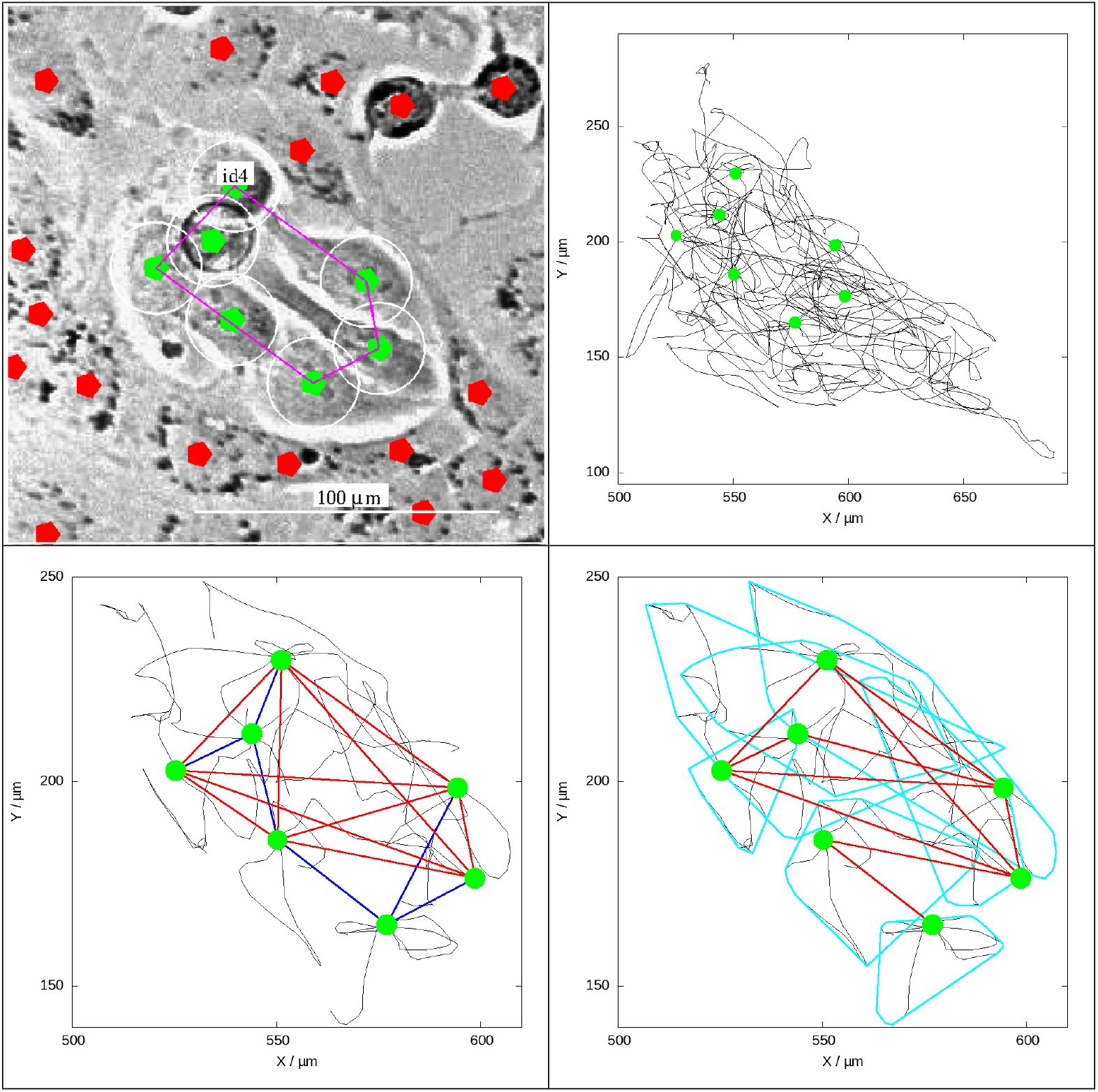
Illustration of two methods, based on call trajectories (black curves), to identify a patch of PC3 cells (green labels) among PC12 cells (red labels).

**Figure 3.**
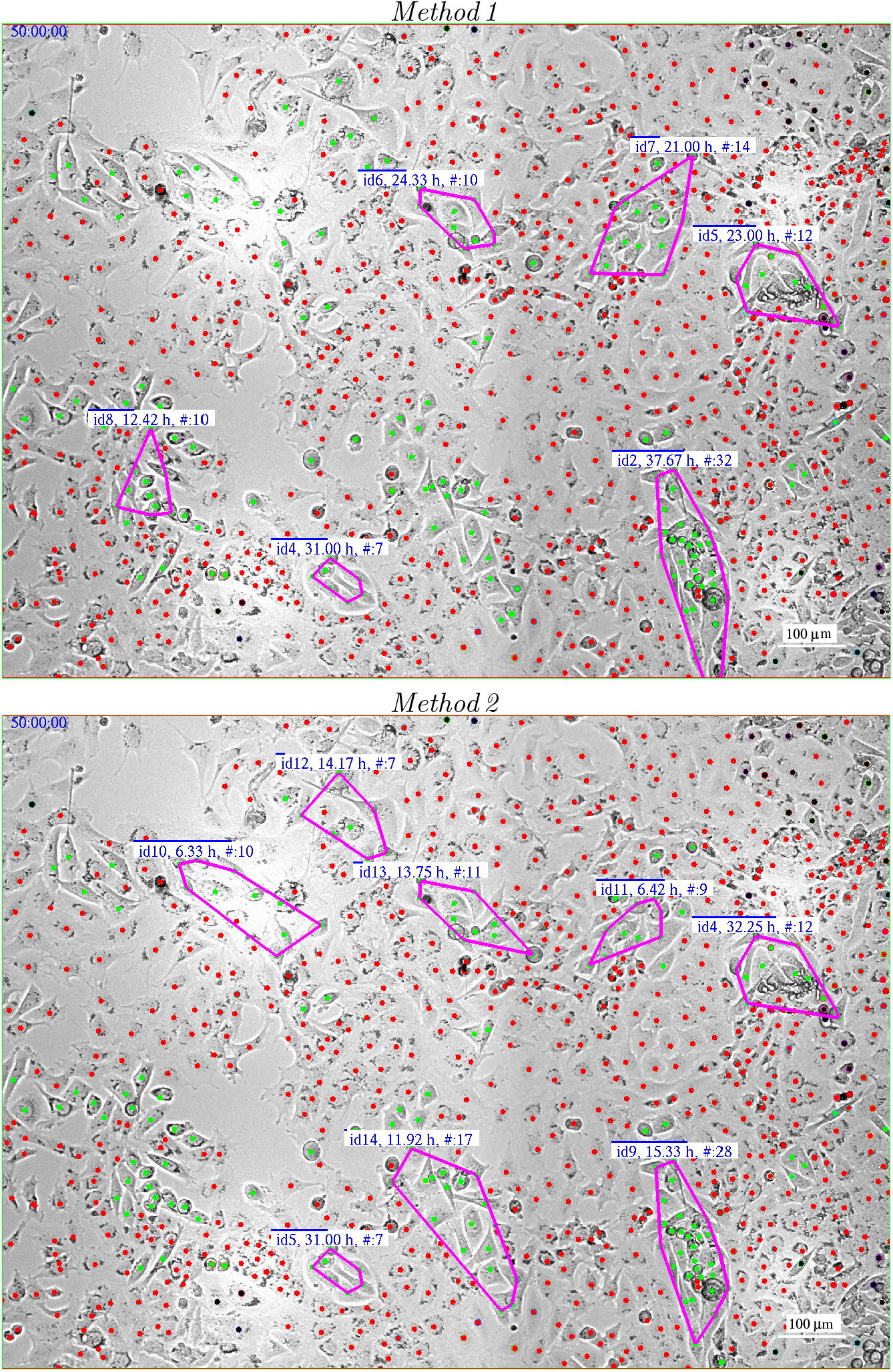
Comparison of *Method 1* and *Method 2* for identifying clustering in PC3 cells (green) in co-culture with PC12 cells (red). The images were captured 50 hours after the start of recording. *Method,2* relies more heavily on cell movement compared to *Method,1*, leading to differences in results, particularly for short-lasting patches. The labels provide the patch ID, duration of the patch, and the current number of cells within the patch.

**Figure 4.**
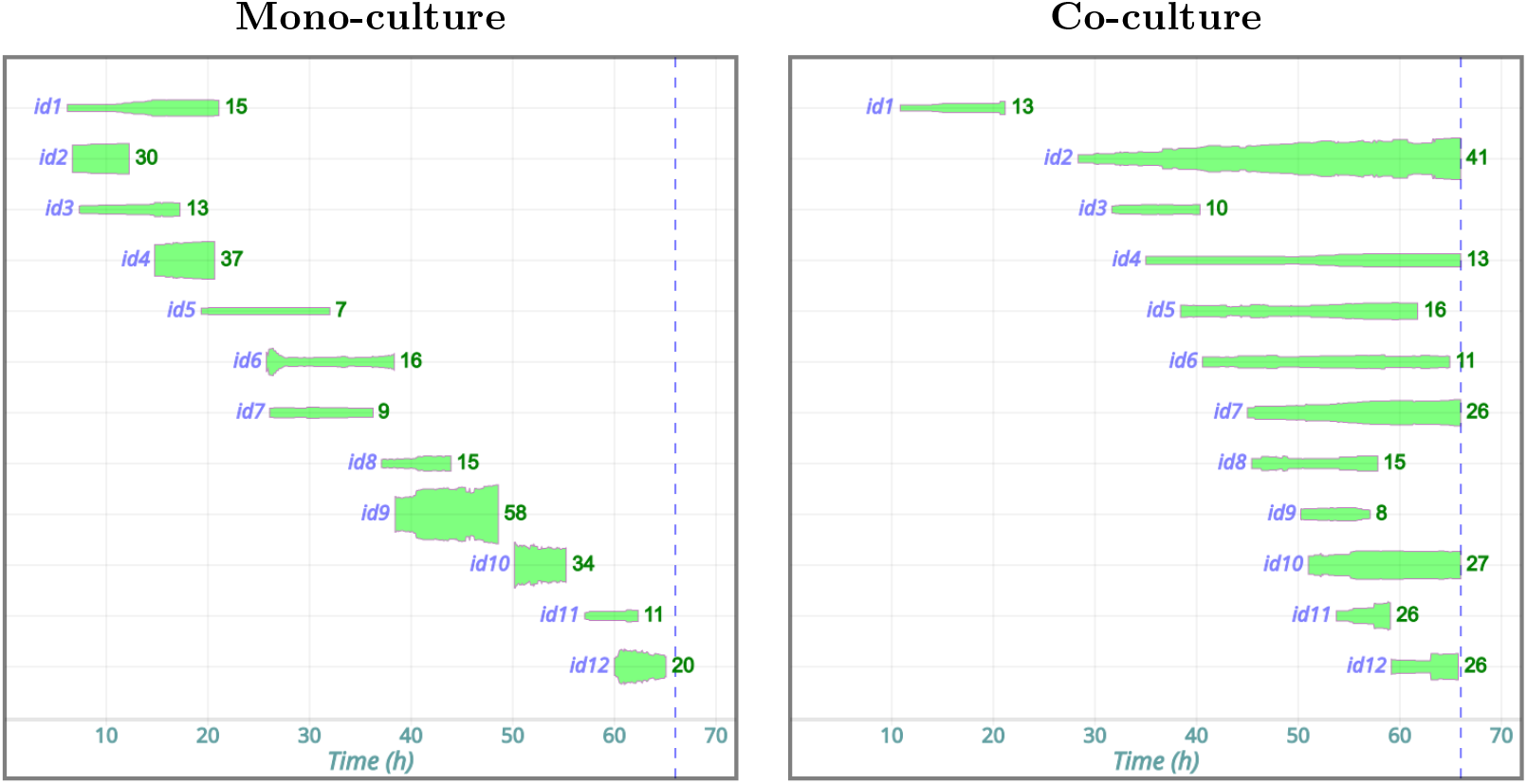
Development of PC3 cell patch sizes in mono-culture and co-culture with PC12 cells, derived from the present experiment data [Korsnes et al., 2024b,c]. Both cultures included 12 patches, identified as *id1, id2*, …, *id12*. The thickness of the elongated green spots represents the number of cells in a patch at a given time after recording began. The numbers to the right of each green spot tells the number of cells at the end of its observation. The blue dotted line indicates the end of the video recording. This figure includes only patches lasting longer than 5 h, which is presumably well above the temporal resolution of the current method for identifying patches (cf. 2.5.3). Notably, patches in the mono-culture appear to dissolve shortly after 10 h. In contrast, many patches in the co-culture persist to the end of recording.

### 2.2 Cell culture

Prostate cancer PC-3 luc2-GFP cells were purchased from Caliper (# 133416) and neuronal PC-12 Adh cell lines were purchased from ATCC (# CRL-1721.1). The PC-3 cells were maintained in RPMI-1640 (# R8758; Sigma) with 10 % FBS (fetal bovine serum)(androgen-proficient medium; F7524; Sigma) and the PC-12 Adh cells were cultivated in F-12K medium ((# 21127022; Thermo Fisher Scientific) with 2.5 % FBS and 15 % horse serum (# 16050122); Thermo Fisher Scientific). Cells were maintained at 37 ^*◦*^C in a humidified 5 % CO_2_ atmosphere. During the cell tracking experiments, both cell lines were maintained in phenol red–free RPMI1640 (# 32404014; Thermo Fisher Scientific) with 10 % FBS. Cells were routinely checked for mycoplasma contamination.

### 2.3 Co-culture establishment

Prostate cancer PC-3 luc2-GFP and PC-12 Adh cells were added in co-culture in a 48 wells plates from Costar ((# 3548). Both cell lines were cultured at a density of 1500 cells per well in RPMI-1640 (# R8758; Sigma) medium without phenol red and 10 % FBS. Cells were then incubated at 37 ^*◦*^C in a humidified 5 % CO_2_ atmosphere for 48 h. Cells in mono-culture were also seeded at a density of 3000 cells per well and incubated using the same conditions as the co-culture. All wells were visually inspected to confirm that there were no abnormalities in the wells and that cells look viable.

### 2.4 Time-lapse video microscopy

PC-12 Adh and PC-3 GFP cells were cultured in mono-or co-culture for time-lapse imaging. Cells were imaged with Cytation5 (BioTek) with temperature and gas control set to 37 ^*◦*^C and 5 % CO_2_ atmosphere, respectively. Sequential imaging of each well was taken using 10 × objective and with an interval of 5 min for 66 h incubation period. Four images with. 5 % overlap from each well were used to stitches together the images used for tracking (Gen5 software; BioTek).

### 2.5 Video-tracking of cells

#### 2.5.1 Cell positional data

Single-cell tracking was performed using the in-house experimental software Kobio Celltrack^1^ for convenience. Alternatively, Fiji Schindelin et al. [2012] and TrackMate Ershov et al. [2022] could also generate the required tracking data. Another viable option is Btrack Ulicna et al. [2021]. The in-house system allows users to define a rectangular region in the center of the video, sized to contain a specified minimum number of cells of each type at the start of recording [Korsnes and Korsnes, 2015, 2018, Quinsgaard et al., 2024]. In the co-culture experiment, a frame was set to include 40 PC3 cells at the start, which automatically included 80 PC12 cells. In the monoculture experiment, the frame was defined to contain 50 PC3 cells. These cells and their descendants were tracked throughout the experiments. Videos demonstrating the performance of the tracking system are available for validation through sample tests Korsnes et al. [2024b,c].

The initial step of the tracking provides cell positional data, which is susceptible to errors that can influence estimates of track length and cell speed. Irregularities in cell movements can inherently make track length scale-dependent or dependent on both temporal and spatial resolution [Korsnes and Korsnes, 2023]. Also note that the current sampling interval of 5 minutes results in missing intermediate positions. Additionally, the precise definition of a cell’s position may not be straightforward. While the centre of the nucleus might be considered a natural reference point, it’s not always discernible in various scenarios, and some cells exhibit multiple nuclei.

The present work takes a pragmatic approach to the above problematic issues and employs a Gaussian filtering of the track positions as follows [Quinsgaard et al., 2024]. Assume that the positional vector ***r***_*i*_ represents the location of a cell at time *t*_*i*_. Define the (Gaussian) weighted sum for all these *n* observed positions of the cell:

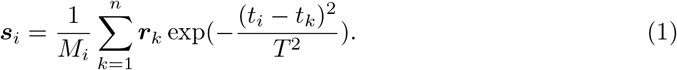

for *i* = 1, 2, …, *n* and where 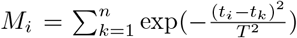. This renders ***s***_*i*_ as a smoothed (filtered) version of the time series ***r***_*i*_ with bandwidth *T*. This work applies *T* = 15 min, considering the sample period is 150 s and that cells tend to move a distance less than their diameter during 15 min. Tests show that final results seem not to be sensitive to perturbations of *T*.

#### 2.5.2 Cell type identification

Video tracking enables the classification of cells into distinct cell types based on single, near-simultaneous GFP fluorescence images, typically acquired at the commencement of the recording. Once a cell’s classification is determined, its entire lineage inherits the same designation. Figure 1 exemplifies this approach.

### Automatic patch identification

This work demonstrates two types of algorithms for identifying clusters of PC3 cells that move together as a unit, even as individual cells within the cluster rearrange positions. The approaches rely on analysing cell movement data over short time periods, limiting the temporal resolution of the methods. These methods, exemplified by Figure 2, differ in how they define which cells are considered ‘linked’ or ‘related’ based on their movement patterns. Graph theory then serves as the basis for the subsequent data treatment, where cells are represented by **vertices** (also known as **nodes**. When two cells are defined to be related or ‘linked’, this relation is called an **edge**. The number of edges (links) of a node *x* is normally written *G*(*x*).

#### *Method 1* : Contact Network Formation

This method defines two PC3 cells as ‘linked’ at time *t* if they come within △*d* = 30 µm of each other within a time frame of △*t* = 2.5 h before and after *t*. A patch (of PC3 cells) is then defined as a collection of nodes (cells) connected via a ‘backbone’ of nodes *x* with degree *G*(*x*) ≥ *G*_min_ = 5. Cells directly linked to this backbone are also considered part of the patch. The backbone network serves to avoid the influence of stray cells on the identification of patches. It also prevents groups of stationary cells from forming patches, as it is unlikely for a cell to have at least five close neighbours simultaneously. Note that the use of a backbone can reduce the ability to observe small clusters of size less than five cells.

#### *Method 2* : Convex Hull Overlap

This method relies on computing the convex hulls of PC3 cell tracks within △*t* = 2.5 h before and after *t*. Two cells are considered linked at time *t* if their respective convex hulls overlap. In this case, the illustrations below do not apply a backbone as in *Method,1*, and a patch is simply defined as an interconnected subset of the linked PC3 cells. This method serves only as an example to illustrate approaches alternative to *Method,1*. It may be of interest as an alternative or complement to *Method,1* since it enables the identification of small clusters.

#### Identity of patches during time

The following outline applies for both methods above. Let ***C***_1_ and ***C***_2_ represent the set of cells in patches at subsequent image times *t*_1_ and *t*_2_, respectively. They are given the same unique identity if

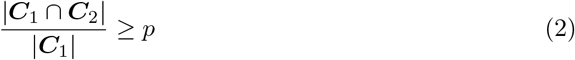

where *p* = 0.75 and |·| represents the size (cardinality) of a set (i.e., the number of cells). Additionally, the mean positions must be within △*r* = 75 µm. In other words, patches retain their identity across subsequent image times if most of the cells at one time step remain in the next, and the patches do not move too much. ID conservation enables determination of duration of patches. This is important to distinguish between behaviours of cells under different treatments.

## 3 Results

### 3.1 Two methods for patch identification

Figure 2 illustrates the above two approaches (*Method 1* and *Method 2*) to identify patches of PC3 cells in co-culture with PC12 cells. These methods differ in that *Method 2* relies more on cell movements compared to its counterpart, *Method 1*, which links two cells at a specific time point, *t*, if they are close together (within a distance △*d* = 30 µm) for a moment within a time window △*t* = 2.5 h before and after *t*. This means cells don’t necessarily need to move much to be linked. However, a cell must link to at least five other cells to become part of the actual network’s backbone, which prevents groups of stagnant cells from forming a patch (see Section 2.5.3). Hence, if cell density makes cells to slow down their movements, *Method 1* is not likely to find patches among them.

*Method 2* uses cell movements to define links between cells. Two cells are considered “linked” at time *t* if the convex hulls of their respective paths intersect during the same time window, as in *Method 1*. This means that the cells must cross each other’s paths to be linked. Therefore, the two methods for patch identification can yield different results (Figure 3).

### 3.2 Comparison of cells in mono-and co-culture

The following data treatment uses *Method 1* for patch identification, leaving *Method 2* as a potential alternative for further work. Patch formation among PC3 cells seems to depend on their environment, with patches appearing to dissolve in mono-culture (Figure 4). Several of these short-lived patches of PC3 cells in mono-culture may be artefacts of the current identification method.

Patches observed among PC3 cells in co-culture appear to persist longer than those seen in monoculture. Notably, the patch labelled *id4* in the co-culture appears around 35 h and remains throughout the entire recording (Figures 4 and 5). Figure 2 shows this same patch at 50 h, along with the tracks of the cells contained within it. These tracks indicate cell-cell cohesion. If the cells were moving independently, the observed track lengths suggest that they would disperse across a much larger area. Hence, the patch cannot be merely an artefact of the patch identification method. Figure 5 further supports this conclusion by depicting the tracks of the initial cells in ***id4*** and their descendants. It also indicates movement of the whole patch.

**Figure 5.**
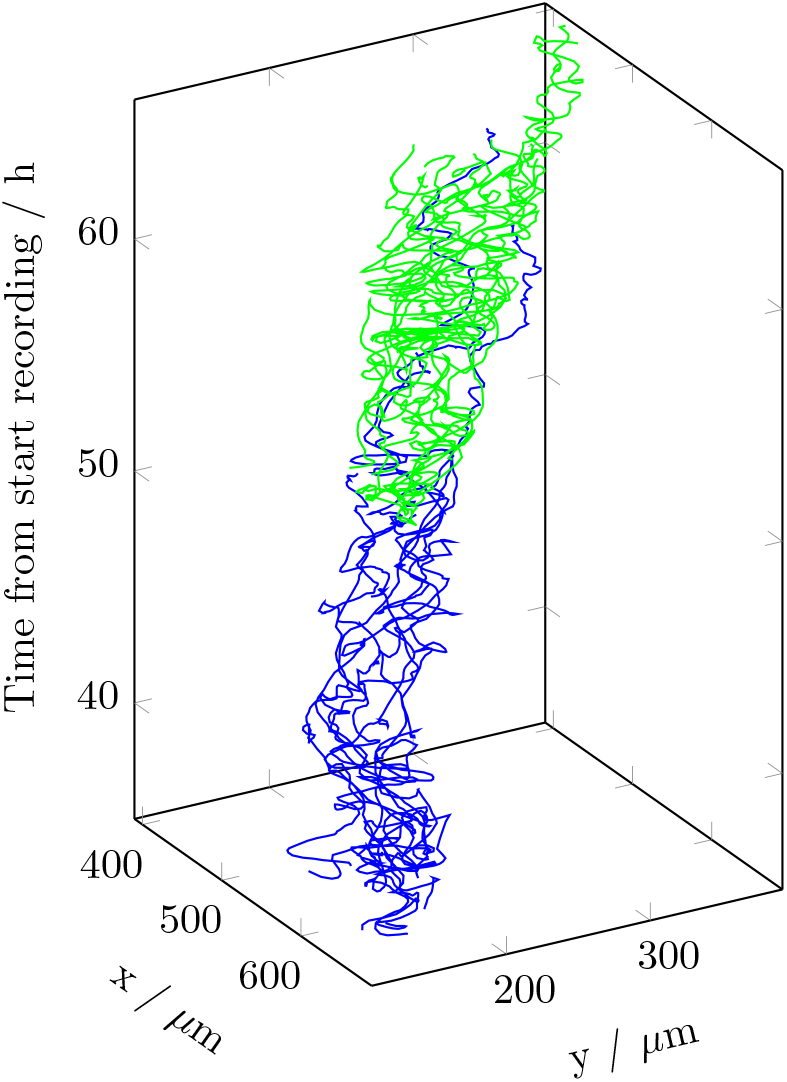
Trajectories of the initial PC3 cells in patch ***id4*** of the co-culture example, including their descendants. Blue and Green represent the first and second generations from the start of the patch, respectively. There is no third generation. The patch appears at 35 h from the start of video recording and persists throughout the recording, which ends at 66 h.

The observed data, visualized in Figure 6, reveals segregation among PC3 cells in mono-culture. Here, groups of cells exhibit distinct movement speeds. This segregation of cells suggests intriguing cellular interactions, deserving further investigation through concepts like flocking or synchronization. Figure 7 further supports the concept of segregation among the PC3 cells. It shows distributions of six-hour average cell speed of PC3 cells in mono-culture and co-culture. Compared to co-culture, PC3 cells in mono-culture exhibit a broader distribution of speeds, indicating a greater heterogeneity in their movement patterns.

**Figure 6.**
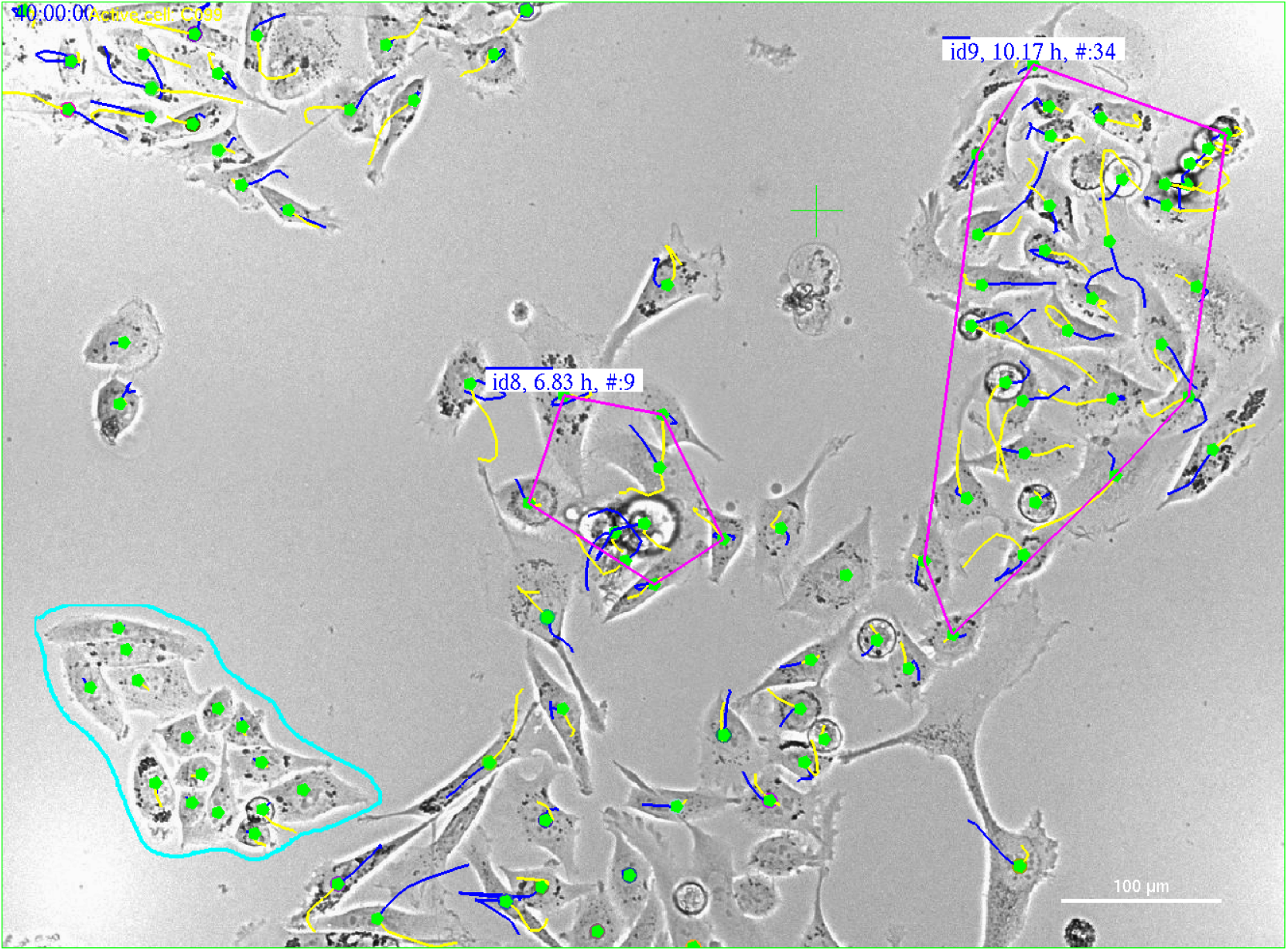
PC3 cells in monoculture: Illustration of differences in cell velocity among groups of cells 40 hours after start of recording. The blue and yellow curves represent the trajectories of cells within 30 min before and after the image time, respectively. Therefore, the length of the tracks indicates individual cell velocities. The cells encircled by the aquamarine curve (in the lower left corner) are significantly more stagnant than those in the patches encircled by magenta lines. As in Figure 3, the labels provide the patch ID, duration of the patch, and the current number of cells within the patch.

**Figure 7.**
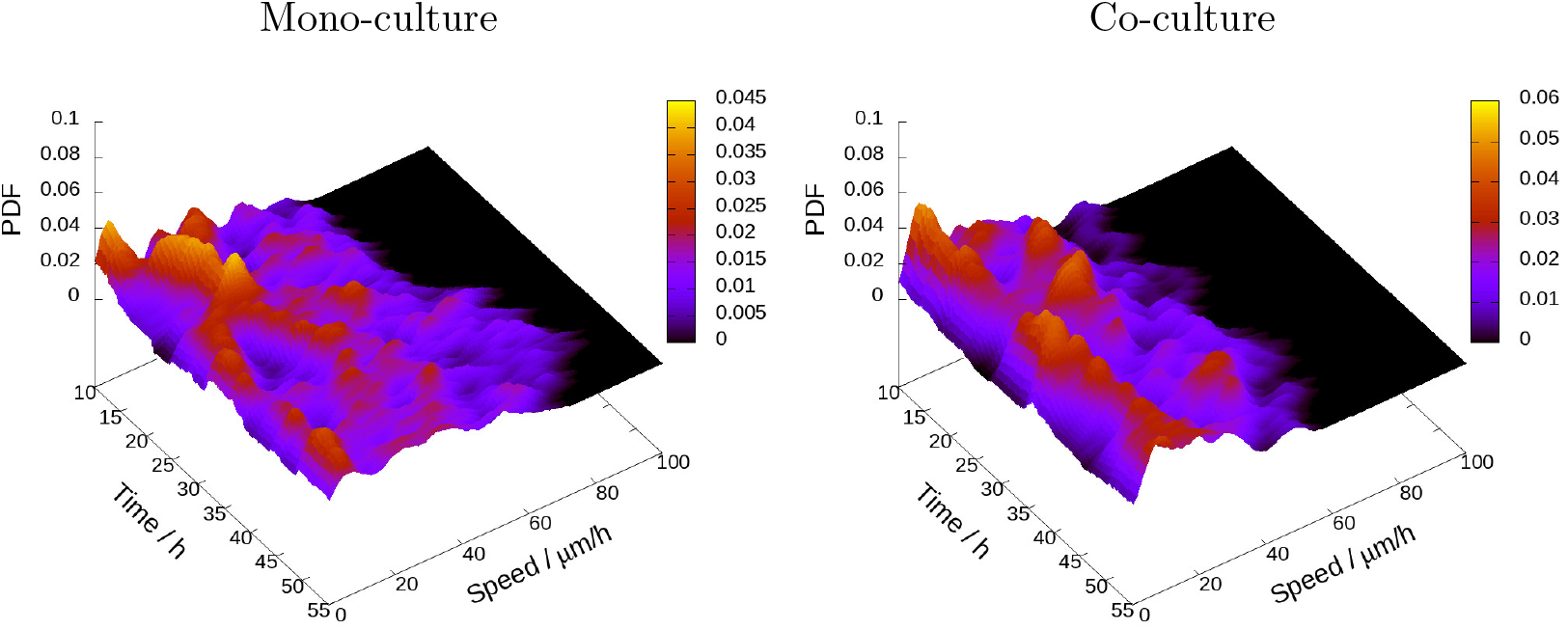
Probability density function (PDF) for six-hour average cell speed during the period from 10 to 55 h after the start of video recording for all tracked PC3 cells in mono-culture (left) and co-culture with PC12 cells (right). The scale bar indicates the colour coding corresponding to different levels of probability density. The cells in mono-culture show a broader variation compared to the cells in co-culture.

A linear increase in the mean squared displacement (MSD) suggests diffusive motion [Huda et al., 2018, Qian et al., 1991, Reynolds, 2018]. Figure 8 builds on this concept and offers a preliminary test for potential cell clustering. It illustrates the temporal evolution of the MSD of PC3 cells in both mono- and co-culture over the period from 40 to 60 hours after recording began. The movement of PC3 cells in mono-culture follows a pattern characteristic of diffusive motion, whereas in co-culture, the cells appear to be more confined

**Figure 8.**
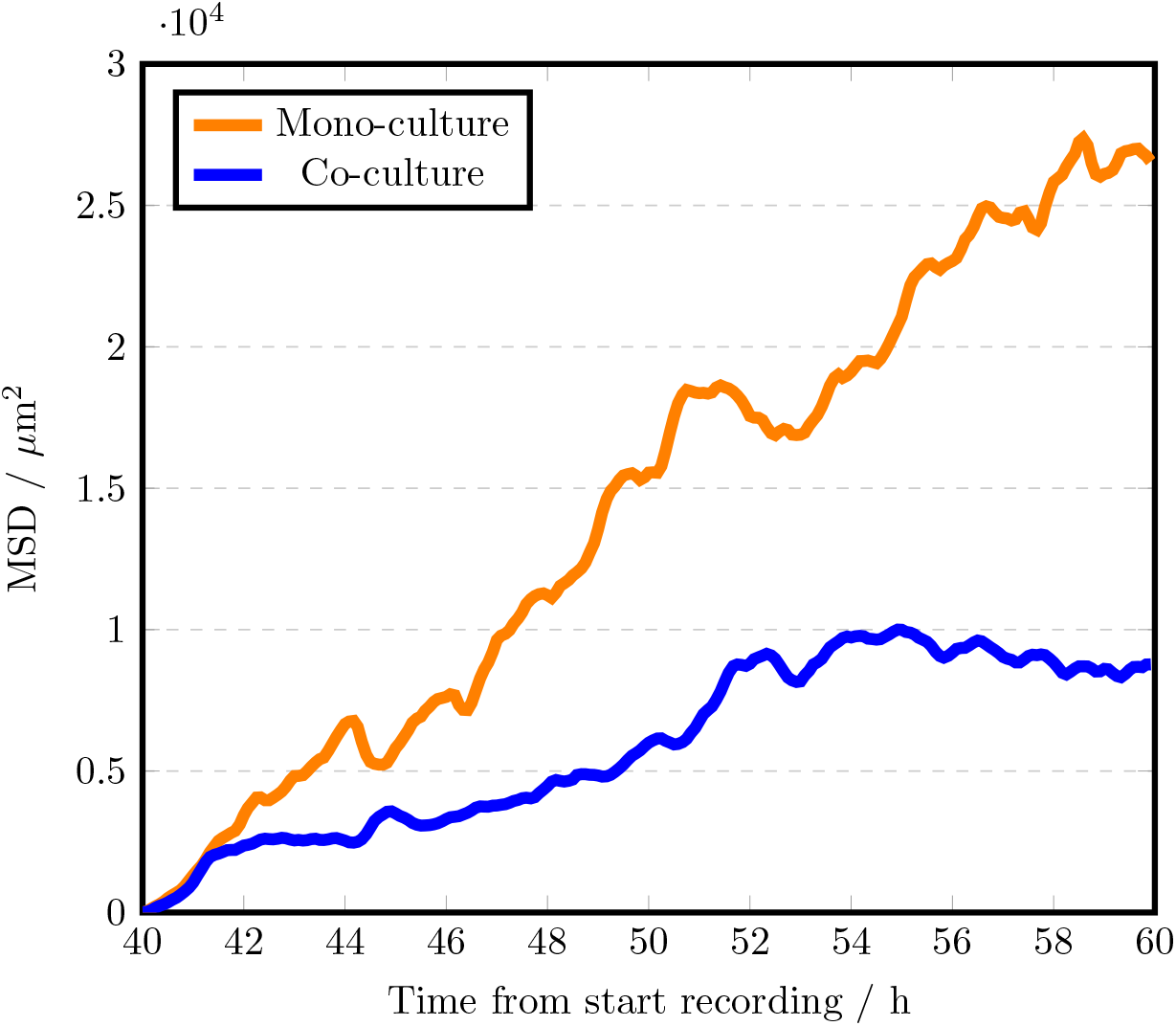
Mean square displacement (MSD) for all complete PC3 cell trajectories between 40 and 60 h from the start of video recording. The orange curve represents PC3 cells in mono-culture (35 trajectories), while the blue curve represents PC3 cells co-cultured with PC12 cells (30 trajectories). Please note that these descriptive statistics are based on a single experiment, where cells in the co-culture appear to follow a particular configuration. Therefore, this data summary does not provide an average across multiple experiments.

- **Upper left:** Subset of video image showing a patch of PC3 cells among PC12 cells at 50 h after the start of recording. The patch is identified as ‘*id4*’ (cf. Figure 4 below).
- **Upper right:** Trajectories of cells in the patch within during their lifetime, starting at 35 h from the beginning of the recording until its end, suggesting a stable cluster.
- **Lower left:** Illustration of *Method 1* for patch identification.

- The black curves here represent trajectories of cells 2.5 h before and after the image time.
- Red and blue lines depict the backbone and peripheral parts of the network, respectively (see Section 2.5.3).
- Backbone nodes (cells) have at least five connections (*G*(*x*) = *G*_min_ ≥ 5), in contrast to others. 8

- **Lower right:** Illustration of *Method 2* where **aquamarine** lines show the convex hull of the tracks of individual cells during 2.5 h before and after the image time (total 5 h). Overlapping convex hulls of two cells’ tracks define a link between them, creating a network (red lines) that defines a patch.

Trajectories of cells forming a patch 35 h after start of recording to clusters. Figure 9 further indicates that cohesion between spatially close PC3 cells affects their movements, leading them to form clusters.

**Figure 9.**
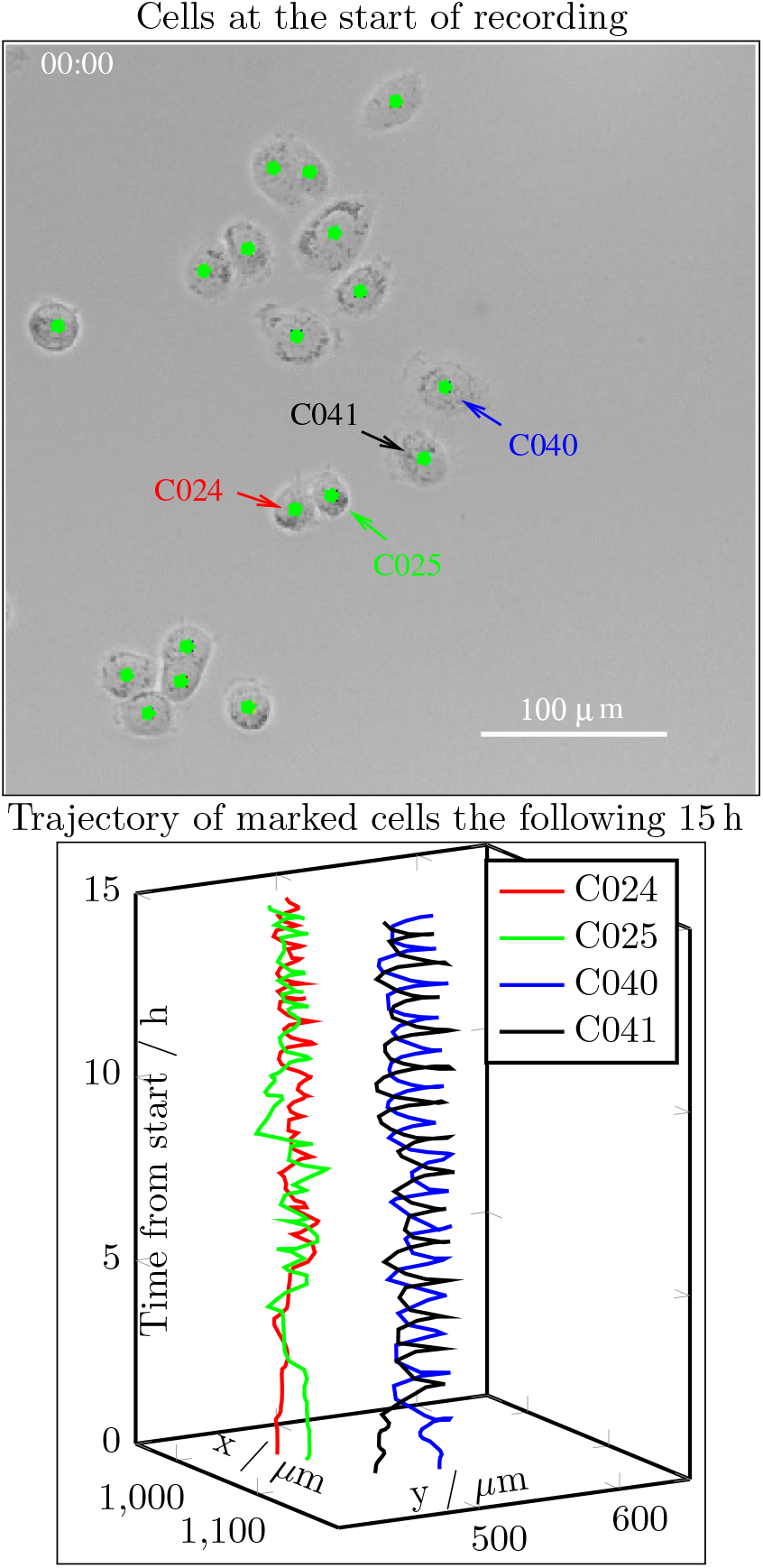
Cell trajectories reveal dynamic interactions between PC3 cells. Left panel: Phase contrast image of PC3 cells in mono-culture at the start of recording. Coloured arrows mark four cells. Right panel: Trajectories of the marked cells over 15 hours, highlighting intercellular interactions. These observations suggest that dynamic cell-cell interactions contribute to the initiation of clustering. See supplementary video [Korsnes et al., 2024a].

Example of mean squared displacement (MSD) of PC3 cells during 20 h

## 4 Discussion

The current data suggest that PC3 cells in co-culture with PC12 cells tend to form lasting clusters. A simple intuition is that cohesion between PC3 cells causes them to expel PC12 cells, leading to the formation of clusters. Subsequently, they may synchronize internal processes and become a distinct and prevailing entity. This presumably depends on the size of the patches. Contact with an environment of similar (PC3) cells may lead to the dissolution of a patch, as indicated by Figure 4. This could prevent further synchronization to initiate. A treatment that affects cell cohesion could help test this hypothesis.

The tendency to form persistent clusters in co-culture provides an opportunity for more in-depth study of clustering. The approach used to identify clusters, inspired by graph clustering theory [Schaeffer, 2007], seems to offer a valuable tool for such investigations. However, developing a generic cluster identification algorithm that can be applied across a wide range of experimental conditions and cell types would require more extensive testing. For example, the current algorithm, *Method 1*, relies on specific values for the parameters △*p*, △*t*, △*d, G*_min_, and *r*. A potential generalization could involve calculating these parameters automatically to identify the most persistent clusters below a certain size, as long-lived small clusters may be particularly relevant to many applications related to cancer spread.

The following equation can replace Equation 2 to define the conservation of patch identity over time:

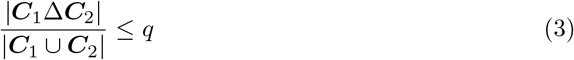

for a value of *q* well below 1, where Δ represents the symmetric set difference (i.e., |***C***_1_Δ***C***_2_| is the number of cells that are in either ***C***_1_ or ***C***_2_ but not in both). This ensures that identified patches do not conserve identity if they suddenly increase in size.

The current video recordings are too short to capture situations where cells slow down due to high confluence, whereas *Method 1* and *Method 2* depend on cell movement. If the cells in a patch or their descendants do not (slowly) disperse, then the patch may be assumed to exist. Notably, if cells within certain clusters have entered a migratory state, one might expect them to be the last to slow down due to increased confluence. This could provide an opportunity for video-based observation of such changes in states.

Theories on clustering graph ^2^ are relevant for a wide range of applications and provide inspiration for generalizing the present approach. Note that a graph consists of vertices (nodes) that are pairwise connected by edges, with associated weights reflecting ‘similarities’ between two vertices (cells) or their spatial distance. A possible modification of *Method 1* is to apply as weight a *Gaussian* kernel:

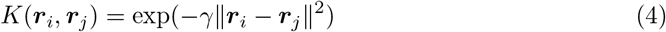

where *γ* = 1*/*(2*σ*^2^) and *σ* determines the width of the kernel [Von Luxburg, 2007]. Here, ***r***_*i*_ and ***r***_*j*_ represent the spatial position of the nodes *i* and *j*.

Several measures of ‘similarities’ between two cells can be combined to reflect membership in a cluster or partition of a graph through cutting wedges based on ‘cost’ of cuts [Nascimento and De Carvalho, 2011]. This approach provides the opportunity to include cellular phenotypes in addition to movements in the process of identifying cellular clusters. The present test data indicate that distributions of cell speed, displacement and clustering parameters differ for PC3 cells in mono- and co-culture (cf Figures 4, 7 and 8). Such distributions may therefore serve as simple general indicators of clustering tendencies and help check for consistency of results from experiments. A more comprehensive data analysis is necessary to definitively incorporate cell speed and morphology into an algorithmic method for cluster identification.

The framework of graph theory allows for the application of theories on percolation and diffusion on graphs [Chung, 1997, Stauffer and Aharony, 2018] to further refine cluster identification from basic tracking data. These theories provide measures of how well a graph or cluster (subgraph) is connected.

An intriguing speculation is that small clusters of closely interacting cells may serve as potential ‘sites of ignition’ for collective changes in behaviour among cancer cells through synchronization, a well-documented phenomenon observed in interacting organisms that influences fundamental cellular processes [Hannon and Ruth, 2014]. The present identification of clustering may therefore represent only an initial step in exploring collective changes among subsets of cells. The current co-culture experimental setup has the potential to facilitate the formation of small clusters, supporting such investigations of collective behaviour. Qualitative studies are a natural initial step in exploring this theory of possible sporadic “sites of ignition”.

A simple, intuitive analogy for this work is that juggling multiple challenges at once is akin to being caught in a ‘fire overlap’ situation, where things become significantly more complex compared to handling each challenge individually. This creates opportunities for cancer cells to migrate as a group. There may be links between cell cluster formation, poor disease prognosis, and resistance to drug treatments [Andrei et al., 2020, Au et al., 2016, Bithi and Vanapalli, 2017, Lee et al., 2017]. Their greater colonization efficiency may stem from factors such as protection against anchorage-dependent apoptosis, cooperation among heterogeneous cell phenotypes within the clusters, and protection from assaults by immune A simple, intuitive analogy forcells [Hong and Zu, 2013, Hou et al., 2012, Yu et al., 2013].

Knowledge of cell cluster biogenesis and organization can contribute to a better understanding of cancer dissemination. Clusters serve as useful monitoring tools and prognostic biomarkers, especially in metastasis, likely due to cell jamming or collective migration mechanisms that remain largely unknown [Amintas et al., 2020]. A promising therapeutic approach suggests that inhibiting or disrupting cluster formation could reduce cancer dissemination [Au et al., 2016, Gkountela et al., 2019].

The present data analysis shows that PC3 cells can exhibit coordinated movement within stable clusters when grown in co-culture with PC12 cells. Cluster sizes from these data align with previous findings of Au et al. [2016], who identified clusters of 20 cells in malignant melanoma. However, clusters of 2 to 6 cells are more commonly reported in the literature [Molnar et al., 2001].

One aspect worthy of investigation is whether cancer cells may collaborate to achieve perineural invasion (PNI), a complex phenomenon characterized by reciprocal interactions between cancer cells and the surrounding nerve micro-environment [Bakst and Wong, 2016, Grigore et al., 2015, Magnon, 2015]. PNI can occur in various malignant tumours [Demir et al., 2012, Schmitd et al., 2018] and it is believed to play a role in the development of neuroendocrine prostate cancer (NEPC), an aggressive variant of prostate cancer where prostate adenocarcinoma cells (PCA) trans-differentiate into a neuroendocrine (NE) cell phenotype to escape anti-androgen therapies [Braadland et al., 2015, Kaarijärvi et al., 2021]. The infiltration of cancer cells into this space can actively stimulate cancer progression and metastasis [Chen et al., 2019, March et al., 2020].

Schwann nerve cells can act as ‘leaders’, reorganizing cancer cell clusters to facilitate PNI when in contact with them [Deborde et al., 2016]. They can also collectively function as tumour-activated Schwann cell tracks (TAST), promoting cancer cell invasion and migration [Deborde et al., 2022]. Physical contact among cells is required to enhance cancer invasion. Cluster detection often correlates with higher rates of disease progression and poorer treatment responses [Wrenn et al., 2021].

It is becoming evident that single-cell profiling allow researchers to address the co-occurrence of molecular events in individual cells, however, it does not need to be limited to gene expression [Kumar et al., 2017]. The incorporation of visualization data on a per-cell basis will complement the knowledge gained from single-cell molecular profiling, including epigenetic modifications, which often contributes to ‘stochastic’ expression [Angermueller et al., 2016].

There have been some limited contributions using in vitro co-cultures to study cell clustering behaviour [Wrenn et al., 2021]. Recent research, however, emphasizes the role of specific molecular mechanisms in promoting cluster formation, which is linked to an increased risk of metastasis. Tumour cells that cluster within the tumour microenvironment (TME) tend to gain an evolutionary advantage. The TME is a complex ecosystem comprising tumour cells, immune cells, fibroblasts, blood vessels, and extracellular matrix (ECM). Clustering facilitates cooperation among tumour cells, enhancing their ability to survive, migrate collectively, spread, and evade the immune system. The presence of cell clustering is often associated with poor prognosis and a heightened likelihood of metastasis in various cancer types [Rozenberg et al., 2023]. Disrupting these mechanisms could offer a promising avenue for cancer treatment, making efforts to uncover them of significant interest.

## 5 Conclusions

PC3 cells cultured in monolayers in test wells, alongside PC12 cells, tend to form stable clusters. This behaviour may be attributed to cohesion between PC3 cells, providing an opportunity to generate data on cell clustering that could be of interest for further studies and may promote new therapeutic ideas. Graph theory offers a framework for developing tools to identify clusters within cell populations. A key step in this process is defining relationships (‘similarities’) between pairs of cells, which creates an initial large graph. This pairing can be based on tracking individual cells over time. The next step involves partitioning the initial large graph into disjoint subgraphs (clusters). A third step is to find similarities between consecutive clusters. This study demonstrates an example of applying this approach.

## Acknowledgments

This study was supported by Olav Raagholt and Gerd Meidel Raagholts legacy, Gidske and Peter Jacob Sørensen research foundation, Astri and Birger Torsteds legacy, Oslo University Hospital and University of Oslo.

## Author Contributions

M.S.K. Conceptualization, Writing original draft, Investigation, Methodology, Data curation, Visualization, Review and editing. H.A.R. Conceptualization, Investigation, Methodology, Data curation, Formal analysis, Visualization, Writing – Review and editing. K.A.T. Conceptualization, Investigation, Writing, Review and editing, Funding acquisition. R.K. Conceptualization, Methodology, Data curation, Formal analysis, Visualization, Software, Writing – Review and editing.

## Ethics statement

Not applicable.

## 6 Data availability

Video illustrations of cell clustering and cell coherence (Figure 9) are available from Mendeley Data [Korsnes et al., 2024a,b,c].

## Footnotes

1 https://www.korsnesbiocomputing.no/

2 https://www.sciencedirect.com/topics/computer-science/clustering-graph

